# A multimodal study of a first episode psychosis cohort: potential markers of antipsychotic treatment resistance

**DOI:** 10.1101/2021.05.03.442450

**Authors:** Kun Yang, Luisa Longo, Zui Narita, Nicola Cascella, Frederick C. Nucifora, Jennifer M. Coughlin, Gerald Nestadt, Thomas W. Sedlak, Marina Mihaljevic, Min Wang, Anshel Kenkare, Anisha Nagpal, Mehk Sethi, Alexandra Kelly, Pasquale Di Carlo, Vidyulata Kamath, Andreia Faria, Peter Barker, Akira Sawa

**Affiliations:** Departments of Psychiatry, Johns Hopkins University School of Medicine, Baltimore, MD 21205, USA; Departments of Neuroscience, Johns Hopkins University School of Medicine, Baltimore, MD 21205, USA; Departments of Biomedical Engineering, Johns Hopkins University School of Medicine, Baltimore, MD 21205, USA; Departments of Genetic Medicine, and Johns Hopkins University School of Medicine, Baltimore, MD 21205, USA; Departments of Radiology and Radiological Science, Johns Hopkins University School of Medicine, Baltimore, MD 21205, USA; Department of Mental Health, Johns Hopkins University Bloomberg School of Public Health, Baltimore, MD 21205, USA; Department of Public Health Studies, Johns Hopkins University Zanvyl Krieger School of Arts and Sciences, Baltimore, MD 21218, USA; Department of Applied Math and Statistics, Johns Hopkins University Whiting School of Engineering, Baltimore, MD 21218, USA; Lieber Institute for Brain Development, Johns Hopkins Medical Campus, Baltimore, MD 21205, USA; F. M. Kirby Research Center for Functional Brain Imaging, Kennedy Krieger Institute, Baltimore, MD 21205, USA

## Abstract

Treatment resistant (TR) psychosis is considered to be a significant cause of disability and functional impairment. Numerous efforts have been made to identify the clinical predictors of TR. However, the exploration of molecular and biological markers is still at an early stage. To understand the TR condition and identify potential molecular and biological markers, we analyzed demographic information, clinical data, structural brain imaging data, and molecular brain imaging data in 7 Tesla magnetic resonance spectroscopy, from a first episode psychosis cohort that includes 138 patients. Age, gender, race, smoking status, duration of illness, and antipsychotic dosages were controlled in the analyses. We found that TR patients had a younger age at onset, more hospitalizations, more severe negative symptoms, a significant reduction in the volumes of the hippocampus (HP) and superior frontal gyrus (SFG), and a significant reduction in glutathione (GSH) levels in the anterior cingulate cortex (ACC), when compared to non-TR patients. The combination of multiple markers provided a better classification between TR and non-TR patients compared to any individual marker. Our study shows that ACC GSH, HP and SFG volumes, and age at onset could potentially be trait biomarkers for TR diagnosis, while hospitalization and negative symptoms could be used to evaluate the progression of the disease. Multimodal cohorts are essential in obtaining a comprehensive understanding of brain disorders.

## Introduction

Psychotic disorder is one of the most debilitating mental conditions, and its disease trajectory is heterogenous [1]. A significant proportion of patients with psychosis continue to have symptoms and poor outcomes despite treatment [2–4]. Clozapine is the only evidence-based medicine for treatment resistant (TR) cases [2], a possible subset of schizophrenia. Clozapine can provide a significant benefit to a substantial portion of TR patients [5]. Nevertheless, due to severe adverse effects [2, 6], clozapine use is still limited in many countries [2]. Paradoxically, it is also known that delayed initiation of clozapine is related to poor response to this medication in TR patients [7, 8]. Thus, identification of patients that are TR, or have susceptibility to TR, at the first clinical visit is very important in order to potentially start clozapine as early as possible. To achieve this goal, it is imperative to establish predictive markers for TR.

In the past decades, multiple groups have put forth major efforts in identifying markers associated with TR. In these studies, the criteria for TR are relatively consistent but also vary depending on the goal of the study. The core criteria for TR patients includes unsuccessful use of two or more non-clozapine antipsychotics and/or the use of clozapine in their past history [2–4, 9, 10]. Thus far, these studies that compare between TR patients and non-TR patients have mainly focused on clinical and demographic markers, and reported that TR patients tend to have a longer duration of untreated psychosis (DUP), more severe negative symptoms, younger age of onset, and poor pre-morbid functioning [4, 11–13]. In addition, some studies have also pointed out severer cognitive deficits in TR patients compared with non-TR patients [14, 15].

Meanwhile, some studies have assessed treatment response during a short interval (for example, 12 weeks) to test whether candidate biological markers at both molecular and anatomy/circuitry levels are associated with treatment response [16–26]. Through such a candidate target approach, increased level of the sum of glutamate and glutamine (Glx) in the anterior cingulate cortex (ACC) estimated through 3-Tesla magnetic resonance spectroscopy (3T-MRS), was reportedly associated with persistent psychotic symptoms despite antipsychotic treatment [18–20]. Whereas, elevated striatal dopamine synthesis capacity was observed in treatment responders measured by positron emission tomography (PET) [21–23]. One group hypothesized that patients with higher glutathione (GSH) levels in the ACC would demonstrate a shorter time to treatment response, and this hypothesis was proven through a 7T-MRS study [24]. Furthermore, other studies have indicated cortical folding defects, white matter regional vulnerability [25], lower cortisol awakening response, and higher interleukin (IL)-6 and IFN-γ for poor treatment response [16, 17].

In the present study, we used a first episode psychosis (FEP) cohort with multimodal datasets, including demographic and clinical information as well as structural and molecular brain imaging data [27–31]. Olfactory functions were also evaluated in this study, since olfactory deficits frequently accompany FEP and may have a predictive value for disease severity and poor prognosis [28, 32, 33]. The goal of this study is to identify markers that can differentiate TR from non-TR patients, in which TR is defined based on medication history [2–4, 9, 11–15], rather than treatment response [16–24]. By taking advantage of multimodal datasets, rather than single layer data, we aimed to obtain a comprehensive landscape associated with TR in psychosis.

## Subjects and Methods

### Study participants

This study was approved by the Johns Hopkins Medicine Institutional Review Board. All study participants provided written informed consent. Patients (n=138) were recruited from within and outside of the Johns Hopkins Hospital. The study psychiatrists did not make treatment decisions regarding medications. The inclusion criteria for all study participants were: 1) between 13 and 35 years old; 2) no history of traumatic brain injury, cancer, abnormal bleeding, viral infection, neurologic disorder, or mental retardation; 3) no drug or alcohol abuse in the past three years; 4) no illicit drug use in the past two months. The inclusion criterion for patients was: at the first visit (baseline), patients must be within 24 months of the onset of psychotic manifestations as assessed by study team psychiatrists using the Structured Clinical Interview for DSM-IV (SCID) and collateral information from available medical records. In the present study, we refer to these patients as FEP patients to differentiate this cohort from recent-onset psychosis cohorts in which patients with a longer duration of illness (e.g., with onset within 5 years) are usually included [34, 35]. We acknowledge that a 2-year-window is a relaxed definition of FEP. Nevertheless, this definition has been used in many published studies and meta-data analyses [36–39]. The majority of patients, except 6 patients, were already medicated at their first (baseline) visit. The 6 medication-naive patients at the baseline started medication during the follow-up period. Patients were diagnosed with schizophrenia (n=72), schizoaffective disorder (n = 14), schizophreniform disorder (n = 3), bipolar disorder with psychotic features (n = 27), major depressive disorder with psychotic features (n = 9), substance induced psychotic disorder (n=5), brief psychotic disorder (n=2), and not otherwise specified psychotic disorder (n=6). Healthy controls (n=115) were also recruited but not used in the present study. After the enrollment (first visit), the study participants were followed up to 4 years.

### Symptomatic and neurocognitive assessment

We used the Scale for the Assessment of Negative Symptoms (SANS) and the Scale for the Assessment of Positive Symptoms (SAPS) to evaluate the presence and severity of negative and positive symptoms respectively [40]. The global and total scores collected at the first (baseline) visit were used in the data analysis. In addition, we used a comprehensive neuropsychological battery we previously developed and have used in multiple publications [27, 29, 41–43]. In brief, we obtained cognitive scores scaled in normally distributed standardized units that covered 6 domains: 1) processing speed (calculated from the combined scores of the Grooved Pegboard test and the Salthouse test); 2) attention/working memory (Digit Span and Brief Attention Memory test); 3) verbal learning and memory (Hopkins Verbal Learning test); 4) visual learning and memory (Brief Visuospatial Memory test); 5) ideational fluency (Ideational Fluency assessment for Word Fluency and Acceptable Designs); and 6) executive functioning (Modified Wisconsin Card Sorting test). These 6 cognitive scores and their average (referred to as the composite score) obtained at the first (baseline) visit were used in the analysis.

### Smell Test

The olfactory functioning of study participants was tested. Specifically, subjects were first administered the Sniffin’ Sticks Odor Identification and Discrimination test to evaluate the ability to identify and discriminate 16 different types of odors. Next, participants were administered 2 odor detection threshold tasks utilizing lyral and citralva as active odorants in a counterbalanced order. The task followed a single reversing staircase, forced-choice format in which individuals were presented 2 vials, one with mineral oil and one containing the active odorant diluted in mineral oil. A total detection threshold score was created that reflected the weakest odor concentration reliably identified as stronger than mineral oil. The detailed procedures were described in our previous publications [28, 31]. In the present study, the data obtained at the study participant’s first visit (baseline) was used for analysis.

### 7T-MRS

Participants were scanned using a 7T scanner (Philips ‘Achieva’, Best, The Netherlands) equipped with a 32-channel head coil (Nova Medical, Wilmington, MA). The ‘LCModel’ software package (Version 6.3-0D) was used to analyze the spectra. Metabolite levels were calculated relative to the water or tCr (total creatine) signal from the same voxel and expressed in institutional units (IU, approximately millimolar) [44]. The detailed protocol for acquiring and processing MRS scans has been described previously[27]. In the present study, we used the data obtained at the first visit (baseline) for glutamate (Glu), gamma-aminobutyric acid (GABA), N-acteylaspartate (NAA), and glutathione (GSH) levels in 5 brain regions: the anterior cingulate cortex (ACC), semiovale (CSO), dorsolateral prefrontal cortex (DLPFC), centrum orbitofrontal cortex (OFR), and thalamus (Thal).

### 3T structural brain imaging: brain volume

T1 high-resolution-weighted images (T1-WI) were obtained from a 3T scanner with the following parameters: sagittal orientation, original matrix 170×170, 256 slices, voxel size 1×1×1.2 mm, TR/TE 6700/3.1 ms. The images were automatically segmented and post-processed through the MRICloud [45] (www.MRICloud.org), as described previously [29]. Briefly, in MRICloud the images were processed through the following steps: 1) orientation and homogeneity correction; 2) two-level brain segmentation (skull-stripping, then whole brain); 3) image mapping based on a sequence of linear, non-linear algorithms, and Large Deformation Diffeomorphic Mapping (LDDMM); 4) multi-atlas labeling fusion (MALF), adjusted by PICSL [46]. In this study, we used the multi-atlas set “Adult22_50yrs_26atlases_M2_V9B” that matched with the demography of our population, through which we collected and analyzed the volumes of 136 brain regions, as defined in the parcellation “level 4” which covered the whole brain (**Table S1**) [47].

### TR patients

Patients who had previously taken more than two non-clozapine antipsychotics and/or were currently taking clozapine were classified as TR. Specifically, we collected medication records and clinical interview notes from the patients’ attending physicians prior to the baseline visit and during the follow-up period (up-to-4-years). Experienced psychiatrists (N.C. and L.L.) carefully reviewed these records to determine how many antipsychotics each patient had tried because of poor response to a medication. If a patient switched antipsychotics due to adverse effects, non-adherence, or other reasons that were independent of his/her response to the medication, the individual was not considered TR. Among 138 patients, 32 satisfied the TR criteria (as a result, 106 patients were non-TR). Eighteen had been TR before their first (baseline) visit and fourteen patients became TR after their first (baseline) visit. The length of follow-up varied among patients. The maximal length of follow-up from disease onset in TR patients was 39 months. In the non-TR group, 70 patients were followed longer than 39 months after disease onset. Some of the remaining 36 patients, though we expect the number to be minor, may turn out to be TR once they are followed longer than 39 months after disease onset. Given this potential caveat, we conducted the analysis using datasets from 32 TR patients and 70 non-TR patients who were followed for at least 39 months after disease onset. In addition, we performed an analysis for 32 TR patients and 106 non-TR patients (all non-TR patients) and disclosed the results in the supplementary document.

### Statistical analysis

R 3.5.1 was used to perform statistical analysis. Group comparisons of demographic and clinical data were calculated using independent t-test for continuous variables, and a chi square test for categorical data. Linear regression with age, gender, race, smoking status, chlorpromazine (CPZ) equivalent dose estimated by the Defined Daily Doses (DDDs) method [48], and duration of illness (DOI) at baseline as covariates was performed when global and total SANS/SAPS, neuropsychological data, olfactory functions, brain volumes, and metabolite levels from 7T MRS were compared between TR patients and non-TR patients. When we assessed neuropsychological data, we also included education as a covariate due to its influence on a subject’s capabilities in interpretation and execution of neuropsychological tests [42]. In the assessment of brain volumes, we also considered handedness in addition to the basic set of confounding factors described above to account for the effects of laterality on brain volumes. A permutation test was performed to evaluate statistical significance. The Benjamini-Hochberg (BH) procedure was used for multiple comparison correction. P values corrected with the BH procedure are presented as q values. The analyte was considered significant if its q value was smaller than 0.05. Finally, we constructed general linear regression models to evaluate the classification performance of individual markers as well as the combination of multiple markers. Leave-one-out cross-validation receiver operating characteristic (ROC) curve and area under the curve (AUC) were used to evaluate performance. The 95% confidence interval of AUC was calculated. Additionally, Akaike information criterion (AIC) was calculated to compare models.

## Results

### Characteristics of the cohort

In the present study, we examined 138 FEP patients, including 32 TR and 106 non-TR patients, but did not use 115 healthy controls for the analysis (see Method section). Using baseline data collected at the first visit, we found that TR patients are younger [mean (TR) = 20.53; mean (non-TR) = 22.76; p value = 5.13E-03], have fewer years of education [mean (TR) = 12.13; mean (non-TR) = 13.24; p value = 0.03], and are less employed (TR = 29.03%; non-TR = 52.31%; p value = 4.76E-02) than non-TR patients (excluding those with short follow-ups) (**Table S2A**). There were no significant differences in gender, race, psychiatric family history, or smoking status. The analysis between TR and all non-TR patients, including those with short follow-ups (see Method section for more detail), led to the consistent conclusion with significant differences in age, education, and employment (**Table S2B**).

### Analysis of clinical data between TR and non-TR patients

Next, we compared the clinical characteristics between TR and non-TR patients (excluding those with short follow-ups). As summarized in **Table 1A**, TR patients were found to be younger at disease onset and have more hospitalizations. DOI and antipsychotic dose was not significantly different between TR and non-TR patients. Next, we compared the SANS/SAPS global and total test scores (**Table 1B**). TR patients had a significantly higher SANS total score and global score in avolition (q-value < 0.05), while no significant differences in either the SAPS total or global scores were observed between TR and non-TR patients. Furthermore, we compared the neuropsychological composite score and sub-domain scores and didn’t observe any significant differences between TR and non-TR patients (**Table 1C**). Similar findings were obtained via comparisons between TR and all non-TR patients (including those with short follow-ups) (**Table S3A, B, C**).

**TABLE 1.**
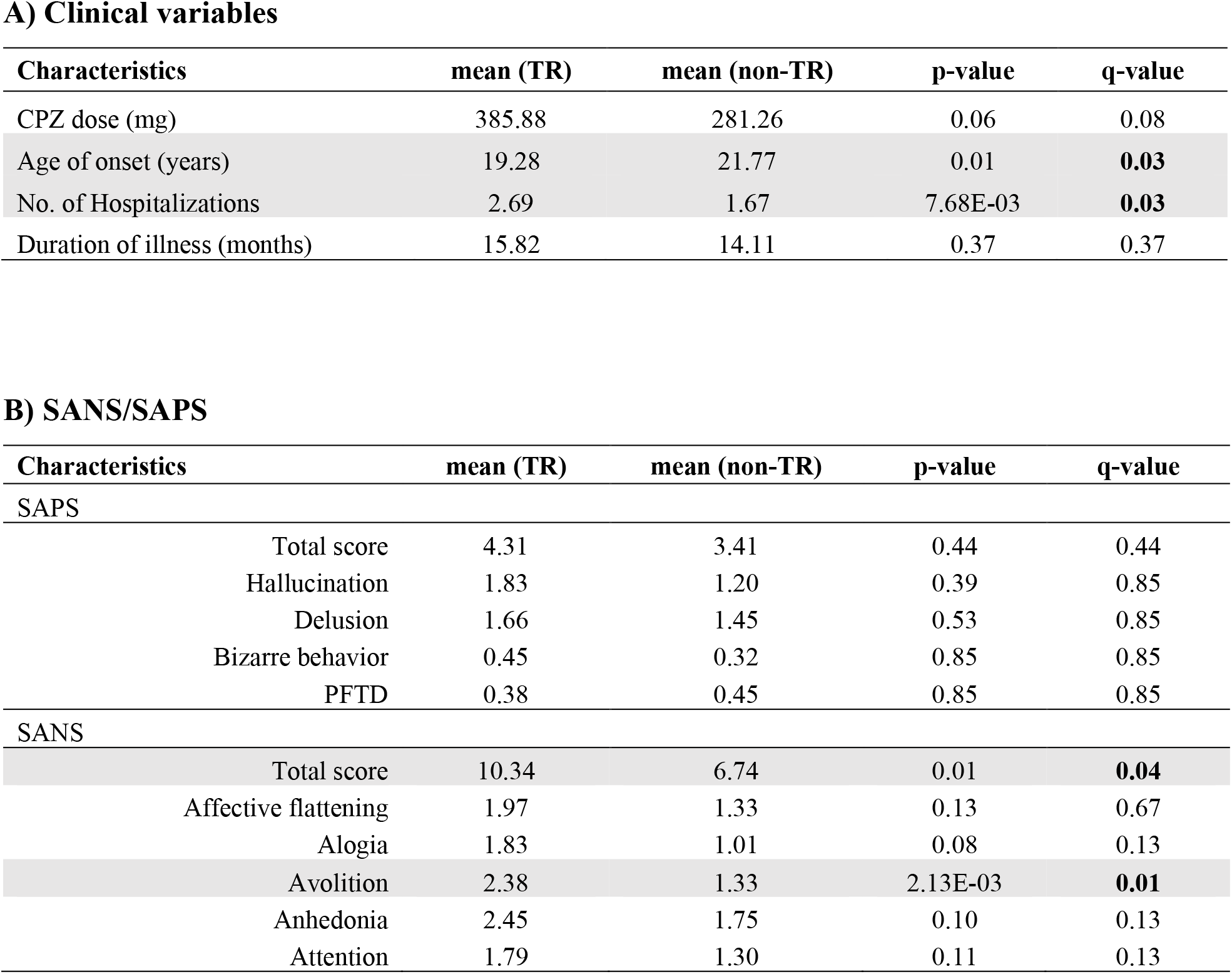
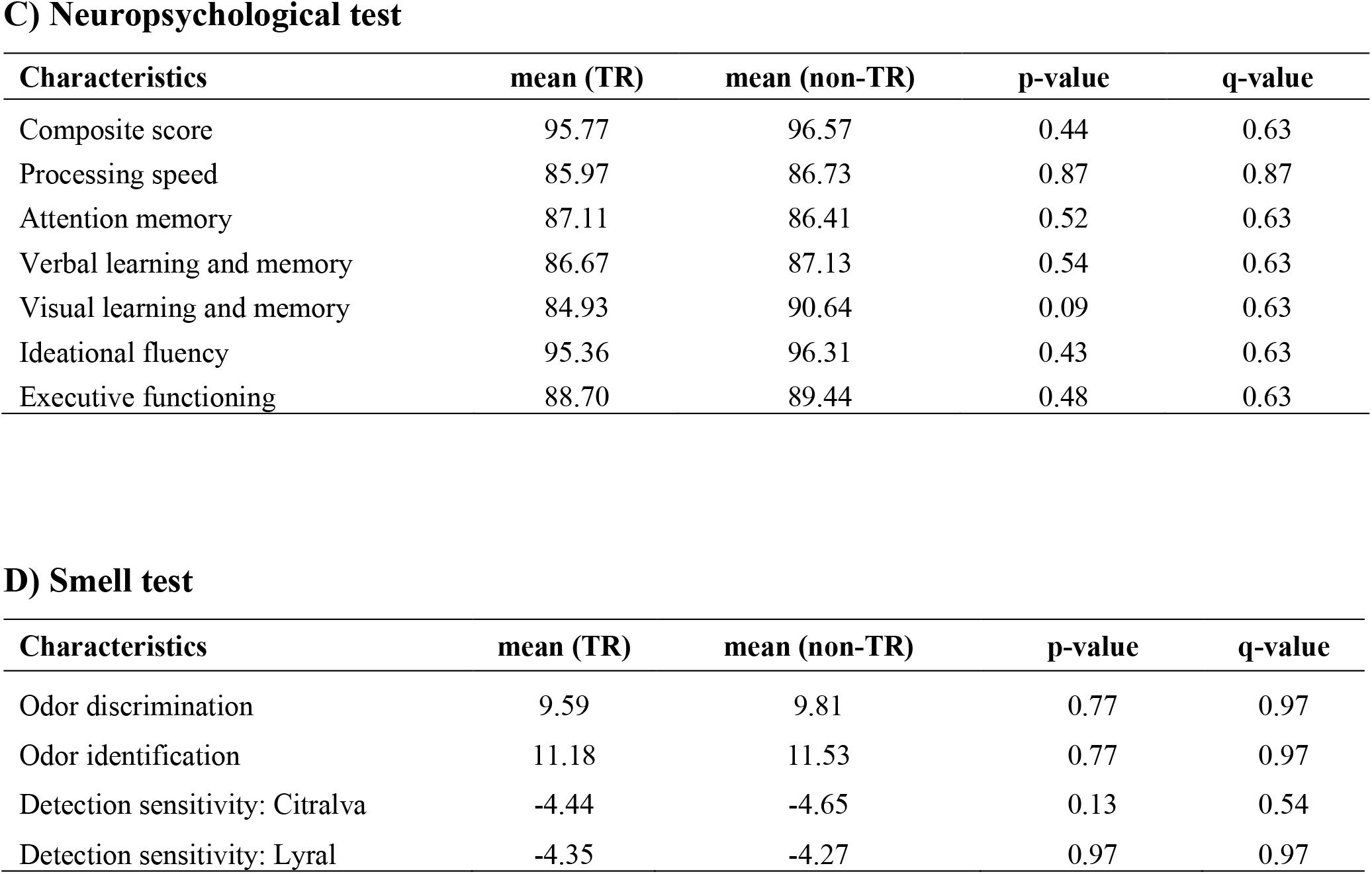
Comparison of clinical data between treatment resistant (TR) and non-TR patients. Linear regression with age, gender, race, and smoking status as covariates was performed to compare clinical data between TR (n=32) and non-TR (n=70) patients. For age of onset, age was not included as a covariate in the linear regression. Significant results (q-value < 0.05) are highlighted in bold with a gray shadow. Abbreviations: SANS represents the scale for the assessment of negative symptoms; SAPS, the scale for the assessment of positive symptoms; and PFTD, positive formal thought disorder. In this table, non-TR patients excluding those with short follow-ups were compared with TR patients.

### Analysis of olfactory test scores between TR and non-TR patients

Previously, our group reported that FEP patients have significantly different odor identification, discrimination, and detection sensitivities of lyral and citralva compared to healthy controls [28]. Thus, we tested whether smell function is different between TR and non-TR patients. We didn’t find significant differences in these olfactory test scores between TR and non-TR patients including or excluding those with short follow-ups (**Table 1D, S3D)**.

### Analysis of brain volume between TR and non-TR patients

Changes in brain volume have been observed in patients with psychosis compared to controls [49–51]. Thus, we wondered whether TR patients have more prominent anatomical changes compared with non-TR patients, and conducted the comparison based on the segmentation in the MRICloud [45]. In this analysis, since no regions met a stringent cutoff of q-values smaller than 0.05, we performed a permutation test to verify the significance of regions with p-values smaller than 0.05. Among 136 studied regions, 35 regions were found to have a significant difference in the volume between TR and non-TR patients (excluding those with short follow-ups) (**Table 2**), whereas 4 regions showed a significant difference in the volume between TR and all non-TR patients (including those with short follow-ups) (**Table S4**). The right hippocampus (HP), left superior frontal gyrus (SFG), left gyrus rectus (GR), and left middle occipital gyrus (MOG) were significantly different between TR and non-TR patients in both comparisons (TR vs. non-TR, excluding or including non-TR patients with short follow-ups).

**TABLE 2.** Analysis results of brain volume between treatment resistant (TR) and non-TR patients. Linear regression with age, gender, race, smoking status, and handedness as covariates was performed to compare brain volume between TR (n=32) and non-TR (n=70) patients. The top 20 brain regions are listed. In this table, non-TR patients excluding those with short follow-ups were compared with TR patients.

| Brain region | Hemisphere | Mean (TR) | Mean (non-TR) | p-value | p-value (permutation test) | q-value |
| --- | --- | --- | --- | --- | --- | --- |
| Middle occipital gyrus | left | 15138.05 | 16462.72 | 4.45E-04 | 0.00 | 0.06 |
| Superior frontal gyrus | left | 24406.95 | 25618.24 | 1.64E-03 | 2.00E-03 | 0.07 |
| Middle frontal gyrus | left | 23062.74 | 24130.76 | 2.59E-03 | 2.00E-03 | 0.07 |
| Occipital white matter | left | 25707.00 | 26786.08 | 2.59E-03 | 2.80E-03 | 0.07 |
| Cingulate | right | 24413.21 | 25674.44 | 3.55E-03 | 0.01 | 0.07 |
| Cuneus | left | 6235.74 | 6656.52 | 3.59E-03 | 2.80E-03 | 0.07 |
| Gyrus rectus | left | 5390.74 | 5734.88 | 3.69E-03 | 0.01 | 0.07 |
| Gyrus rectus | right | 5478.47 | 5724.36 | 4.39E-03 | 0.01 | 0.07 |
| Cingulate white matter | right | 4790.47 | 4962.84 | 4.96E-03 | 3.20E-03 | 0.07 |
| Cuneus | right | 5639.63 | 5975.64 | 0.01 | 0.01 | 0.08 |
| Middle frontal gyrus | right | 22359.63 | 23029.00 | 0.01 | 0.01 | 0.08 |
| Frontal white matter | left | 49371.00 | 50835.32 | 0.01 | 0.01 | 0.08 |
| Occipital sulcus | left | 1038.95 | 1158.92 | 0.01 | 0.01 | 0.08 |
| Cingulum | right | 3182.32 | 3534.16 | 0.01 | 0.01 | 0.08 |
| Superior frontal gyrus | right | 23067.89 | 23849.60 | 0.01 | 0.01 | 0.09 |
| Frontal white matter | right | 51468.32 | 52633.36 | 0.01 | 0.01 | 0.09 |
| Superior temporal gyrus | right | 18287.05 | 18763.04 | 0.01 | 0.01 | 0.11 |
| Hippocampus | right | 3608.11 | 3806.56 | 0.01 | 0.02 | 0.11 |
| Limbic | right | 1894.00 | 2000.00 | 0.02 | 0.01 | 0.11 |
| Lingual gyrus | right | 7173.63 | 7554.24 | 0.02 | 0.01 | 0.11 |

### Analysis of 7T-MRS data between TR and non-TR patients

We previously studied 5 distinct brain regions (ACC, DLPFC, OFR, CSO, and Thal), and reported that FEP patients have significantly different levels of Glu, GABA, NAA, and GSH, compared with healthy controls [27]. With the hypothesis that some of these brain metabolites may be associated with TR, in the present study, we compared these metabolites in all 5 brain regions between TR and non-TR patients. We found that GSH in the ACC was the only significant metabolite (water reference: q-value = 0.03): GSH levels in the other brain regions and the other metabolite levels in all 5 brain regions were not significantly different between TR and non-TR patients (excluding non-TR patients with short follow-ups) (**Figure 1, Table S5**).

**FIGURE 1.**
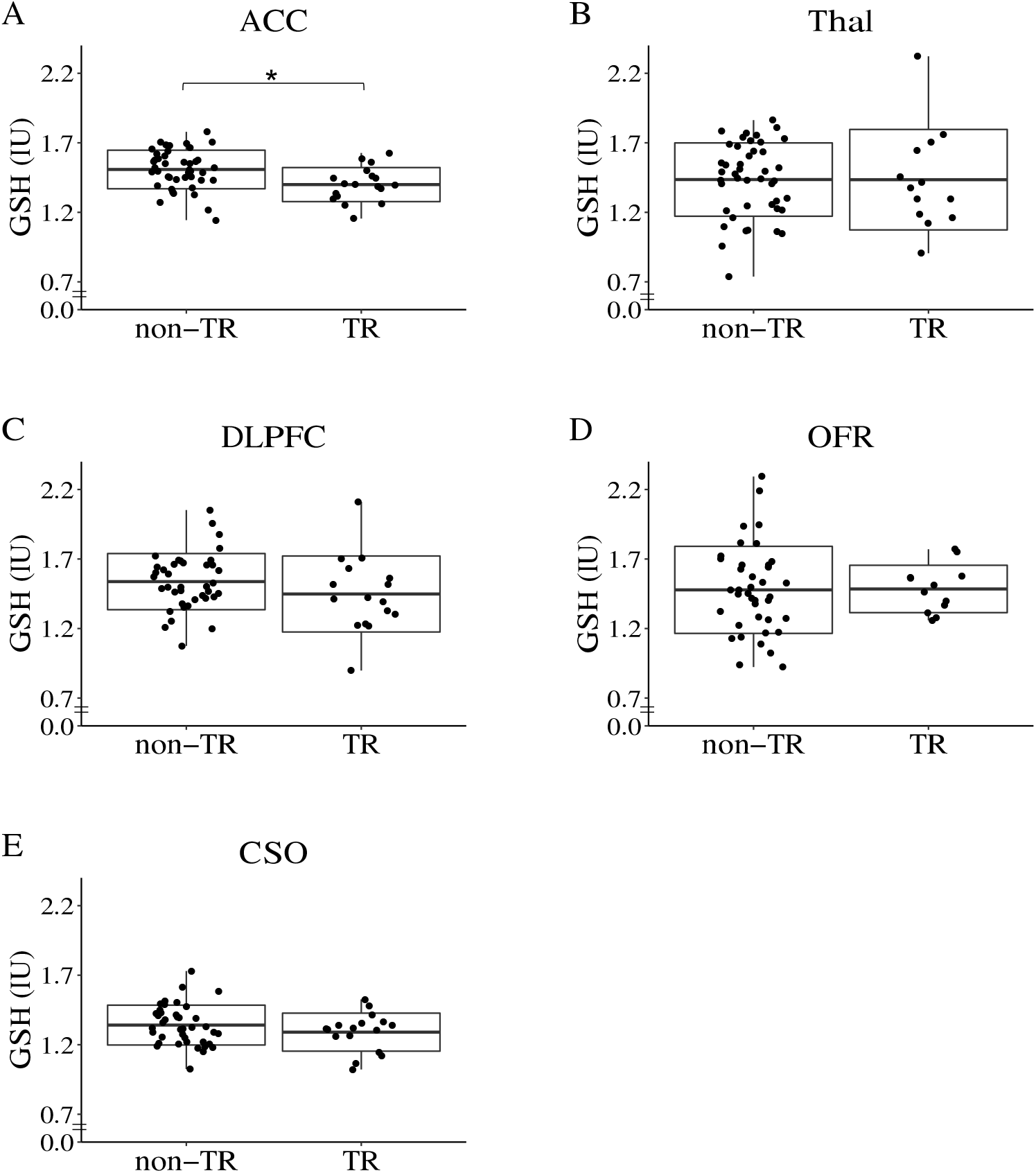
Box plots of glutathione (GSH) levels (water reference) in treatment resistant (TR) and non-TR patients. The box represents standard deviation and the solid line in the middle of the box shows the mean value. Black dots represent individual subjects. Symbol * denotes significant results (q-value < 0.05). Abbreviations: ACC represents anterior cingulate cortex; Thal, thalamus; OFR, orbital frontal cortex; DLPFC, dorsolateral prefrontal cortex; CSO, centrum semiovale, and IU, institutional unit, approximately millimolar. In this analysis, non-TR patients excluding those with short follow-ups were compared with TR patients.

Similar findings were obtained via comparison between TR and all non-TR patients (including non-TR patients with short follow-ups) (**Table S5**).

### Multimodal data for TR and non-TR classification

One unique aspect of the present study is that multimodal data was collected from this FEP cohort, which gives us an opportunity to integrate different types of data to obtain a comprehensive view for TR. Thus far, at least from each individual modality, we observed that TR and non-TR groups are different in age of onset, number of hospitalizations, negative symptoms, the level of GSH in the ACC (ACC-GSH), and volume changes in specific brain regions, in particular the right hippocampus (**Table 3**).

**TABLE 3.** Summary of significant findings. This table summarizes the q-values for CPZ dose, number of hospitalizations, age of onset, ACC GSH, and SANS scores, the range of q-values for multiple scores from smell test, neuropsychological test, and SAPS, and p-values for right hippocampus, left superior frontal gyrus, left gyrus rectus, and left middle occipital gyrus. Stable markers for TR are highlighted in gray shadow. Abbreviations: TR represents treatment resistant; CPZ, chlorpromazine; ACC, anterior cingulate cortex; GSH, glutathione; SANS, the scale for the assessment of negative symptoms; and SAPS, the scale for the assessment of positive symptoms.

|  | TR vs non-TR<br>(short follow-ups excluded) | TR vs non-TR<br>(short follow-ups included) | Change over<br>time |
| --- | --- | --- | --- |
| CPZ dose | 0.08 | 0.04 | 2.86E-03 |
| No. of Hospitalizations | 0.03 | 0.01 | 0.08 |
| Age of onset | 0.03 | 0.01 | 1.00 |
| ACC GSH | 0.03 | 4.64E-02 | 0.92 |
| Right hippocampus | 0.01 | 0.01 | 0.77 |
| Left superior frontal gyrus | 1.64E-03 | 0.02 | 0.47 |
| Left gyrus rectus | 3.69E-03 | 0.04 | 0.01 |
| Left middle occipital gyrus | 4.45E-04 | 0.04 | 0.01 |
| Smell test | > 0.05 | > 0.05 | < 0.05 |
| Neuropsychological test | > 0.05 | > 0.05 | < 0.05 |
| SAPS: total/global score | > 0.05 | > 0.05 | < 0.05 |
| SANS: avolition | 0.01 | 0.03 | 0.03 |
| SANS: total score | 0.04 | 0.03 | 0.11 |

We next asked whether multimodal datasets could provide a better understanding of TR that could differentiate TR patients from non-TR patients. Markers that differentiate these two groups may include state indicators that change over time. On the other hand, trait markers whose values are stable over the disease course or at least in early disease stages are useful to predict the risk or biological vulnerability to TR. Thus, in the present study, we paid attention to trait markers for TR (markers that were shared among patients who met the TR criteria before and after the baseline visit) rather than signatures associated with the TR state. Under such premise, we used classification models with trait markers to evaluate the performance of individual markers as well as the combination of them. Our previous longitudinal assessment for the same cohort used in the present study showed that the ACC-GSH levels were stable in patients between 15-35 years of age [52]. Using the same analytic pipeline that we used to assess longitudinal changes in the ACC-GSH level [52], we found that the right HP volume and the left SFG volume were also stable in FEP patients (**Table S6**: see also details in the supplementary methods). Additionally, age of onset won’t change over time. The SANS total score and the number of hospitalizations were not considered trait markers under the cutoff (q-value > 0.3) that we defined in a conservative manner. Altogether, we identified 4 stable trait markers (ACC-GSH, right HP volume, left SFG volume, and age of onset), which were further evaluated using classification models.

Interestingly, using TR patients and non-TR patients excluding those with short follow-ups, we found that the integration of markers could largely improve the classification performance with an AUC of 0.84, while the AUCs for age of onset, ACC-GSH, right HP, and left SFG were 0.65, 0.70, 0.56, and 0.58, respectively (**Figure 2, Table S7**). The AICs for models using age of onset, ACC-GSH, right HP, left SFG, and all the four markers are 53.20, 51.17, 56.36, 56.27, and 37.59, respectively. Similarly, analysis using TR and all non-TR patients (including those with short follow-ups) found that the combination of multimodal markers outperformed individual markers (**Figure S1, Table S7**). This implies that multimodal datasets could provide complementary or non-redundant information to reach a better understanding of TR. Without a single factor highly associated with TR, a combination of biomarkers may complement the small effect size of individual factors and help us to identify TR patients at an early stage.

**FIGURE 2.**
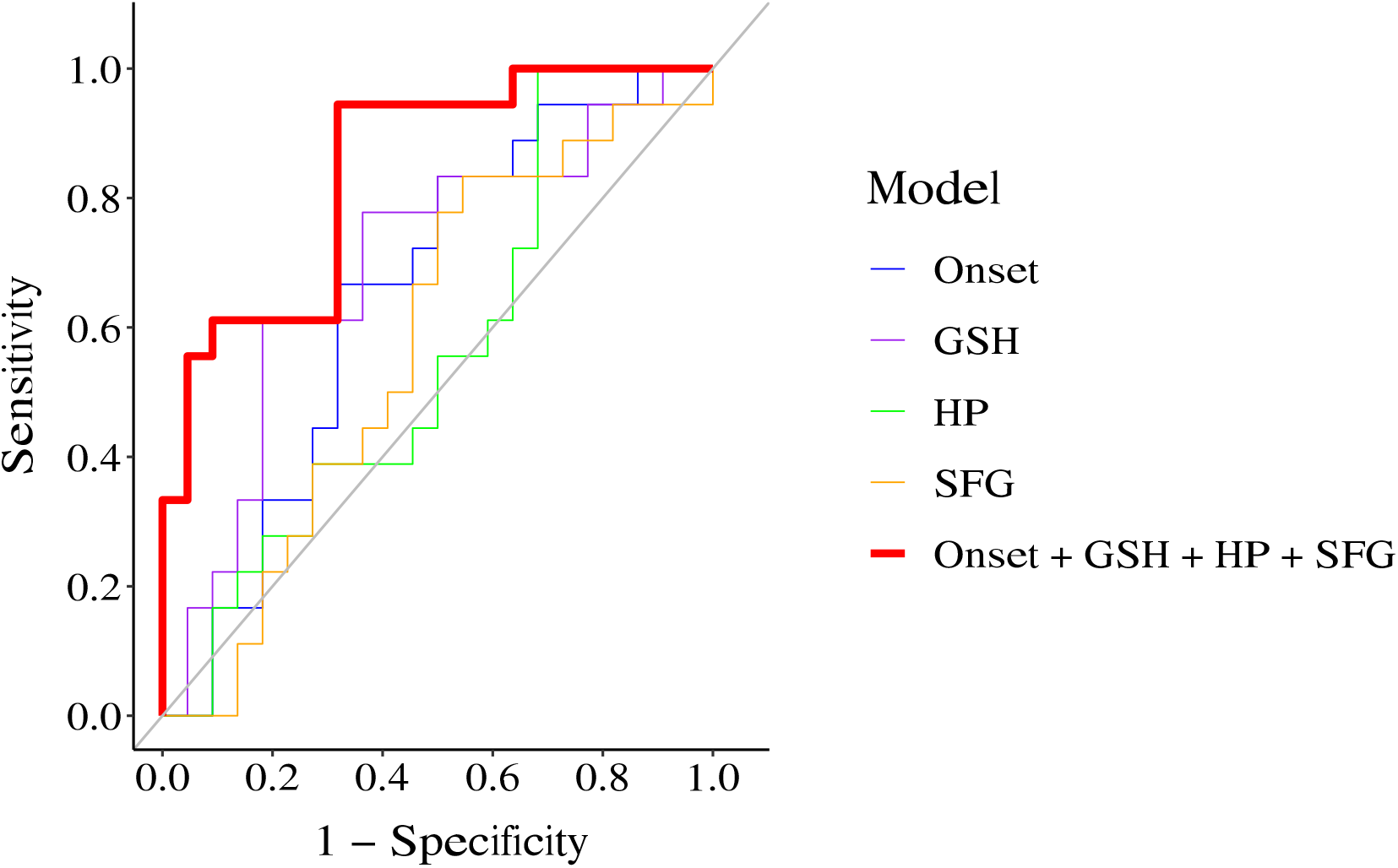
Classification models of treatment resistant (TR) and non-TR patients. We compared the performance of five classification models: 1) model of age of onset (denoted as Onset), blue, AUC=0.65; 2) model of glutathione in anterior cingulate cortex (denoted as GSH), purple line, AUC=0.70; 3) model of right hippocampal volume (denoted as HP), green line, AUC=0.56; 4) model of left superior frontal gyral volume (denoted as SFG), orange line, AUC=0.58; and 5) model of multimodal markers (denoted as Onset + GSH + HP + SFG), red line, AUC=0.84. In this modeling, the data from non-TR patients excluding those with short follow-ups and TR patients were used.

## Discussion

We have established a FEP cohort with multimodal data collection. The percentage of TR patients was consistent with previous findings that about 20-30% of patients develop TR [2–4, 53]. We confirmed that younger age of onset, more hospitalizations, and severer negative symptoms were associated with TR patients when compared with non-TR patients, which is also consistent with previous reports from multiple groups [2–4]. Notably, we showed that GSH was selectively reduced among several key metabolites such as Glu, GABA, and NAA in the ACC in TR patients. Furthermore, a change in GSH was observed only in the ACC with no difference in the levels of GSH in any other brain regions (DLPFC, Thal, CSO, OFR) we studied. Furthermore, by using an unbiased approach using MRICloud [45], we underscored the volume changes that are associated with TR, in particular the right HP and the left SFG.

One unique aspect of this study is that multimodal data was obtained from the same subjects. By taking advantage of this, we constructed classification models with multimodal data that were stable in the trajectory of early disease phases, and demonstrated that a combination of multiple markers led to much better classification performance than individual markers. This suggests that using multiple markers may overcome the issue of small effect sizes of individual markers and identify TR patients at early stages. The performance of the classification models in the present study may not directly impact clinical practice, nevertheless, the goal of this study is to explore the direction of early diagnosis and intervention for TR. Diagnostic strategies using multimodal data have achieved success in the clinical practice of many diseases. For example, a variety of invasive and non-invasive tests including physiological, imaging, and molecular markers are routinely used for the diagnosis of cardiovascular disease [54]. With additional molecular markers from future studies at the epigenetic, gene, or protein levels, we believe that a clinical-grade diagnostic tool employing multimodal data could be developed for TR.

In this study, we used medication records to define TR. Instead of closely tracking the treatment response to antipsychotics during a short interval (such as half to one year), we chose this approach because medication records can be easily collected. Furthermore, this strategy is easily expanded globally beyond potential differences involving hospitalization that tends to be affected by social systems. For example, discoveries from this study can be further validated by population studies or consortia research like ENIGMA (Enhancing Neuro Imaging Genetics through Meta-Analysis) [55]. Nevertheless, it is important that study psychiatrists carefully determine that the switches of antipsychotics are not because of adverse effects, non-adherence, or for other unrelated reasons. Our biological observations that were made in a comprehensive manner (semi-unbiased or unbiased approaches) are compatible with the biological changes that were reported in a candidate target approach for antipsychotic treatment response [24]. These observations suggest that, although study designs based on antipsychotic treatment response for a short period and those that stand on medication records are somewhat disconnected in the overall TR research field, both of them have room to be interpreted in an integrative and comprehensive manner.

In the present study, we observed that TR patients had a significantly higher SANS total score and global score in avolition, whereas no significant differences in either the SAPS total or global scores were observed between TR and non-TR patients. Given that TR is defined as poor response to antipsychotics in regard to psychosis, these results may sound paradoxical. Nevertheless, thus far, more than one meta-data analysis has consistently reported that negative symptoms rather than positive symptoms were significantly different between TR and non-TR groups [56, 57].

A reduction in the HP volume or a decrease in the SFG volume have been reported in patients with psychotic disorder [58–61]. Here, we observed a significant volume reduction in both the HP and SFG in TR patients compared with non-TR patients, making the observation more specific. Furthermore, we also observed a significant reduction in the level of GSH in the ACC. GSH is a physiological reservoir for Glu through the GSH cycle [62], and a reduction in the ACC GSH level, as shown in the present study, implies not only redox imbalance but also glutamatergic neurotransmission imbalance [63] involving the ACC in TR patients. The brain regions underscored in the present study mediate multiple higher brain functions, including associative learning and memory, in which glutamatergic and non-dopaminergic neurotransmission play important roles [64, 65]. This may be one of the reasons why D2-targeted antipsychotics, regardless of first or second generation, may not work for TR patients. We previously reported gray-matter abnormalities in deficit schizophrenia, including major changes in the SFG and ACC [66]. A recent report pointed out the connectivity between the HP and ACC as a potential marker to predict treatment response to second generation antipsychotics in FEP patients [26]. These reports may be interpreted in the same mechanistic context as what we have observed in the present study.

The present study may have potential limitations. First, this study does not include a replication cohort. However, the goal of this study is to discover new directions by using a relatively unique cohort in which multimodal data are available for FEP patients. For this reason, we believe that the inclusion of a replication cohort is beyond the scope of the present work. Nevertheless, the main message is to extract multimodal markers for TR that can be easily collected in many institutions and countries. Among these markers, the MRS data for ACC-GSH may be difficult for some hospitals in some countries. To overcome this dilemma, efforts to identify a surrogate blood marker that correlates with ACC-GSH may be important. In addition, we acknowledge that some of the 106 patients classified as non-TR, might become TR if they were followed longer. To address this possibility, we performed analysis using non-TR patients with and without excluding those with short follow-ups. The results and conclusions of the two-group comparisons were in essence the same. Thus, this dilemma should not affect the overall conclusion. Furthermore, in the present study, the majority of the patients were medicated. Statistical analysis can’t fully address the effects of antipsychotics on brain structures and metabolites. Further studies with medication-naïve patients are expected to confirm our findings. Lastly, we used general linear regression models to evaluate the performance of multimodal markers, which may not be the most efficient way for data fusion. Nevertheless, we employed this approach in the present study, as this is straightforward and can be easily verified by other cohorts.

The research presented here demonstrates the utility and advantages of multimodal study cohorts for psychiatric research. By utilizing deep phenotyping data covering genetics, metabolomics, proteomics, brain imaging, and clinical tests, we can obtain a holistic view of complex and heterogeneous diseases, like psychosis, through hypothesis and/or data-driven approaches (**Figure S2**). We expect that well-maintained multimodal study cohorts will become essential resources for tackling the diagnosis and treatment of brain disorders.

## Acknowledgements

This study is supported by National Institutes of Mental Health Grants MH-092443 (to AS), MH-094268 (to AS), MH-105660 (to AS), and MH-107730 (to AS); foundation grants from Stanley (to AS), RUSK/S-R (to AS), and a NARSAD young investigator award from Brain and Behavior Research Foundation (to AS, KY). The original recruitment of study participants was partly funded by Mitsubishi Tanabe Pharma Corporation. The authors thank Drs. Brian Caffo for kindly contributing to scientific discussions and feedback related to this work. The authors appreciate Ms. Yukiko Lema for research management and manuscript organization, and thank Dr. Melissa A Landek-Salgado for critical reading of the manuscript.

