## Supplementary Materials for "A multimodal study of a first episode psychosis cohort: potential markers of antipsychotic treatment resistance"

[g]Department of Public Health Studies

### SUPPLEMENTARY METHODS

#### Longitudinal analysis of brain volume

Four significant brain regions were identified by comparisons between TR and non-TR (including or excluding those with short follow-ups) patients: left gyrus rectus, right hippocampus, left middle occipital gyrus, and left superior frontal gyrus. We further analyzed the longitudinal changes in volume to see if they could be trait markers. Among the 138 first episode psychosis (FEP) patients recruited in this study, 88 patients came back for annual 3 tesla (T) magnetic resonance imaging (MRI) scans during a 4-year follow-up period. We compared the baseline volumes of these 4 brain regions between the entire cohort (n=138) and the longitudinal cohort (n=88), and didn't find significant differences. This suggests that there is no return bias in the studied cohort that could affect the longitudinal analysis results. Next, we used the analytic pipeline developed by our previous study to analyze the longitudinal changes in brain volume [1]. Specifically, to quantitatively analyze longitudinal changes, we calculated the annual percentage change (APC) by using the following equation:

$$APC = \frac{1}{n-1}\sum_{i=1}^{n-1}\frac{V_{i+1} - V_i}{(D_{i+1} - D_i)/365}$$

$V$ is the level of metabolite, $D$ is the date of the visit, $i$ is the ith visit, $D_{i+1} - D_i$ is the number of days between two visits, and $n$ is the total number of visits. We performed one sample t-test and Bayesian test to test whether the APC was significantly different from zero in FEP patients. The Benjamini-Hochberg (BH) procedure was used for multiple comparison correction. P values corrected with the BH procedure are presented as q values. The analyte was considered significant if its q value was smaller than 0.05. We found a significant decline in the left gyrus rectus and left middle occipital gyrus, while the APC of the right hippocampus and left superior frontal gyrus were close to 0 (**Table S6**).

### SUPPLEMENTARY TABLES

#### TABLE S1. Brain regions analyzed in the present study.

Volumes of the left and right hemisphere of the following regions were used in the present study

| | | **Symbol** | **Full Name** |
|---|---|---|---|
| Gray matter | Frontal | SFG | Superior frontal gyrus |
| | | MFG | Middle frontal gyrus |
| | | IFG | Inferior frontal gyrus |
| | | OG | supra orbital gyrus |
| | | RG | Gyrus rectus |
| | | PrCG | Precentral gyrus |
| | Parital | PoCG | Postcentral gyrus |
| | | SPG | Superior parietal gyrus |
| | | SMG | Supramarginal gyrus |
| | | AG | Angular gyrus |
| | | PrCu | Pre-cuneus |
| | Occipital | Cu | Cuneus |
| | | LG | Lingual gyrus |
| | | SOG | Superior occipital gyrus |
| | | IOG | Inferior occipital gyrus |
| | Temporal | FuG | Fusiform gyrus |
| | | MOG | Middle occipital gyrus |
| | | STG | Superior temporal gyrus |
| | | MTG | Middle temporal gyrus |
| | | ITG | Inferior temporal gyrus |
| | | Insula | Insula |
| | Limbic | Hippo | Hippocampus |
| | | CGH | Hippocampus cingulum |
| | | Amyg | Amygdala |
| | | CGC | Cingulate cortex |
| | | Cingulate | Cingulate gyrus |
| | Deep gray matter | Caud | Caudate nucleus |
| | | Caudate_tail | Caudate tail |
| | | GP | Globus pallidus |
| | | Put | Putamen |
| | | Thalamus | Thalamus |

(continued on next page)

| | | | |
|---|---|---|---|
| White matter | Deep white matter | ALIC | Anterior limb of internal capsule |
| | | PLIC | Posterior limb of internal capsule |
| | | BCC | Body of corpus callosum |
| | | SCC | Splenium of corpus callosum |
| | | GCC | Genu of corpus callosum |
| | | Fx | Fornix |
| | | Fx.ST | Stria terminalis of fornix |
| | | ant_DPWM | Anterior deep white matter (part f corona radiata) |
| | | inf_DPWM | Inferior deep white matter (part o corona radiata) |
| | | post_DPWM | Posterior deep white matter (part f corona radiata) |
| | | PVA_anterior | Anterior periventricular white matter |
| | | PVA_posterior | Posterior periventricular white matter |
| | Peripheral | PeripheralCingulateWM | Cingulate white matter |
| | | PeripheralFrontalWM | Frontal white matter |
| | | PeripheralOccipitalWM | Occipital white matter |
| | | PeripheralParietalWM | Parietal white matter |
| | | PeripheralTemporalWM | Temporal white matter |
| Mesencephalon | | BasalForebrain | Basal forebrain |
| | | midbrain | Midbrain |
| Metencephalon | | Pons | Pons |
| | | CerebellumWM | Cerebellum white matter |
| | | Cerebellum | Cerebellum |
| CSF spaces | Ventricles | III_ventricle | Third ventricle |
| | | AnteriorLateralVentricle | Anterior (frontal) horn of lateral ventricle |
| | | InferiorLateralVentricle | Inferior (temporal) horn of the lateral ventricle |
| | | PosteriorLateralVentricle | Posterior (occipital) horn of lateral ventricle |
| | | IV_ventricle | Fourth ventricle |
| | Sulci | CentralSul | Central sulcus |
| | | CinguSul | Cingulate sulcus |
| | | FrontSul | Frontal sulcus |
| | | OcciptSul | Occipital sulcus |
| | | ParietSul | Parietal sulcus |
| | | TempSul | Temporal sulcus |
| | | SylFrontSul | Frontal pars of sylvian sulcus |
| | | SylParieSul | Parietal pars of sylvian sulcus |
| | | SylTempSul | Temporal pars of sylvian sulcus |

**TABLE S2. Demographic summary of patients.**

Significant results (p-value < 0.05) are highlighted in bold with a gray shadow. Abbreviations: sd represents standard deviation; FEP, first episode psychosis; and TR, treatment resistant.

##### A. TR and non-TR patients (excluding those with short follow-ups)

| Characteristics | TR (32) | non-TR (70) | p-value |
|---|---|---|---|
| Age (mean years ± sd) | 20.53 ± 2.97 | 22.76 ± 4.78 | **5.13E-03** |
| Gender (male/female) | 25/7 | 49/21 | 0.48 |
| Race (black/white) | 16/16 | 35/35 | 1.00 |
| Psychiatric family history (positive/negative/unknown) | 21/8/3 | 45/21/4 | 0.81 |
| Smoking status (yes/no) | 10/22 | 18/52 | 0.63 |
| Years of education (mean ± sd) | 12.13 ± 2.20 | 13.24± 2.69 | **0.03** |
| Employment (yes/no) | 9/22 | 34/31 | **4.76E-02** |

##### B. TR and all non-TR patients (including those with short follow-ups)

| Characteristics | TR (32) | non-TR (106) | p-value |
|---|---|---|---|
| Age (mean years ± sd) | 20.53 ± 2.97 | 22.86 ± 4.72 | **1.28E-03** |
| Gender (male/female) | 25/7 | 77/29 | 0.65 |
| Race (black/white) | 16/16 | 54/52 | 1.00 |
| Psychiatric family history (positive/negative/unknown) | 21/8/3 | 72/28/6 | 1.00 |
| Smoking status (yes/no) | 10/22 | 30/76 | 0.82 |
| Years of education (mean ± sd) | 12.13 ± 2.20 | 13.12 ± 2.42 | **0.03** |
| Employment (yes/no) | 9/22 | 47/46 | **0.04** |

**TABLE S3. Comparison of clinical data between treatment resistant (TR) and all non-TR patients (including those with short follow-ups).**

Linear regression with age, gender, race, and smoking status as covariates was performed to compare clinical data between TR (n=32) and all non-TR (n=106) patients. For age of onset, age was not included as a covariate in the linear regression. Significant results (q-value < 0.05) are highlighted in bold with a gray shadow. Abbreviations: SANS represents the scale for the assessment of negative symptoms; SAPS, the scale for the assessment of positive symptoms; and PFTD, positive formal thought disorder.

##### A) Clinical variables

| Characteristics | mean (TR) | mean (non-TR) | p-value | q-value |
|---|---|---|---|---|
| CPZ dose (mg) | 385.88 | 271.32 | 0.03 | **0.04** |
| Age of onset (years) | 19.28 | 21.95 | 4.17E-03 | **0.01** |
| # Hospitalizations | 2.69 | 1.60 | 1.57E-03 | **0.01** |
| Duration of illness (months) | 15.82 | 13.69 | 0.28 | 0.28 |

##### B) SANS/SAPS

| Characteristics | mean (TR) | mean (non-TR) | p-value | q-value |
|---|---|---|---|---|
| SAPS | | | | |
| Total score | 4.31 | 3.46 | 0.64 | 0.64 |
| Hallucination | 1.83 | 1.23 | 0.59 | 0.81 |
| Delusion | 1.66 | 1.44 | 0.55 | 0.81 |
| Bizarre behavior | 0.45 | 0.34 | 0.81 | 0.81 |
| PFTD | 0.38 | 0.46 | 0.69 | 0.81 |
| SANS | | | | |
| Total score | 10.34 | 7.16 | 0.01 | **0.03** |
| Affective flattening | 1.97 | 1.38 | 0.09 | 0.43 |
| Alogia | 1.83 | 1.11 | 0.14 | 0.16 |
| Avolition | 2.38 | 1.51 | 4.19E-03 | **0.03** |
| Anhedonia | 2.45 | 1.78 | 0.03 | 0.07 |
| Attention | 1.79 | 1.39 | 0.12 | 0.16 |

(continued on the next page)

#### C) Neuropsychological test

| Characteristics | mean (TR) | mean (non-TR) | p-value | q-value |
|---|---|---|---|---|
| Composite score | 95.77 | 95.19 | 0.62 | 0.94 |
| Processing speed | 85.97 | 86.00 | 0.97 | 0.97 |
| Attention memory | 87.11 | 85.09 | 0.49 | 0.94 |
| Verbal learning and memory | 86.67 | 85.32 | 0.86 | 0.97 |
| Visual learning and memory | 84.93 | 87.35 | 0.36 | 0.94 |
| Ideational fluency | 95.36 | 94.57 | 0.67 | 0.94 |
| Executive functioning | 88.70 | 89.10 | 0.52 | 0.94 |

#### D) Smell test

| Characteristics | mean (TR) | mean (non-TR) | p-value | q-value |
|---|---|---|---|---|
| Odor discrimination | 9.59 | 9.70 | 0.63 | 0.86 |
| Odor identification | 11.18 | 11.39 | 0.80 | 0.86 |
| Detection sensitivity: Citralva | -4.44 | -4.58 | 0.19 | 0.76 |
| Detection sensitivity: Lyral | -4.35 | -4.28 | 0.86 | 0.86 |

**TABLE S4. Analysis results of brain volume between treatment resistant (TR) and all non-TR patients (including those with short follow-ups).**

Linear regression with age, gender, race, smoking status, and handedness as covariates was performed to compare brain volume between TR (n=32) and all non-TR (n=106) patients. The top 10 brain regions are listed. Significant results (p-values smaller than 0.05 in both linear regression and permutation test) are highlighted in bold with a gray shadow.

| Brain region | Hemisphere | Mean (TR) | Mean (non-TR) | p-value | p-value (permutation test) | q-value |
|---|---|---|---|---|---|---|
| Hippocampus | right | 3608.11 | 3900.80 | **0.01** | **8.00E-04** | 0.88 |
| Superior frontal gyrus | left | 24406.95 | 25624.78 | **0.02** | **0.02** | 0.88 |
| Gyrus rectus | left | 5390.74 | 5729.19 | **0.04** | **0.02** | 0.88 |
| Middle occipital gyrus | left | 15138.05 | 16108.09 | **0.04** | **0.04** | 0.88 |
| Cuneus | right | 5639.63 | 5928.63 | 0.05 | 0.16 | 0.88 |
| Middle frontal gyrus | right | 22359.63 | 23141.25 | 0.06 | 0.12 | 0.88 |
| Limbic | right | 1894.00 | 1978.14 | 0.08 | 0.04 | 0.88 |
| Syllabus parietal sulcus | left | 206.63 | 258.20 | 0.08 | 0.05 | 0.88 |
| Cuneus | left | 6235.74 | 6563.66 | 0.08 | 0.10 | 0.88 |
| Middle frontal gyrus | left | 23062.74 | 23913.16 | 0.08 | 0.04 | 0.88 |

**TABLE S5. Analysis of the 7T MRS data.**

Linear regression with age, gender, race, and smoking status as covariates was performed to compare the 7T MRS data (normalized by water or total creatine (tCr)) between TR (n=32) and non-TR (n=70) patients (excluding those with short follow-ups), as well as, between TR (n=32) and all non-TR (n=106) patients (including those with short follow-ups). Results with a p value smaller than 0.05 are highlighted in bold with a gray shadow. Abbreviations: Abbreviations: TR represents treatment resistant; GABA, γ-aminobutyric acid; Glu, glutamate; GSH, glutathione; NAA, N-acetylaspartate; ACC indicates anterior cingulate cortex; Thal, thalamus; DLPFC, dorsolateral prefrontal cortex; CSO, centrum semiovale; and OFR, orbital frontal region.

**A. TR and non-TR patients (excluding those with short follow-ups); metabolite levels are normalized by water.**

| Brain region | Metabolite | mean (TR) | mean (non-TR) | p-value | q-value |
|---|---|---|---|---|---|
| ACC | **GSH** | 1.40 | 1.51 | **1.54E-03** | **0.03** |
| | NAA | 7.06 | 7.19 | 0.36 | 0.96 |
| | GABA | 1.71 | 1.68 | 0.58 | 0.96 |
| | Glu | 8.06 | 8.05 | 0.98 | 0.98 |
| CSO | GSH | 1.29 | 1.34 | 0.47 | 0.96 |
| | NAA | 8.32 | 8.26 | 0.24 | 0.95 |
| | GABA | 1.68 | 1.82 | 0.01 | 0.12 |
| | Glu | 5.99 | 6.13 | 0.10 | 0.70 |
| DLPFC | GSH | 1.45 | 1.54 | 0.57 | 0.96 |
| | NAA | 7.98 | 8.34 | 0.28 | 0.95 |
| | GABA | 1.59 | 1.55 | 0.68 | 0.97 |
| | Glu | 6.83 | 7.33 | 0.14 | 0.71 |
| OFR | GSH | 1.48 | 1.50 | 0.83 | 0.98 |
| | NAA | 7.87 | 7.72 | 0.53 | 0.96 |
| | GABA | 1.67 | 1.68 | 0.98 | 0.98 |
| | Glu | 6.53 | 6.69 | 0.44 | 0.96 |
| Thal | GSH | 1.43 | 1.44 | 0.78 | 0.98 |
| | NAA | 7.43 | 7.60 | 0.63 | 0.97 |
| | GABA | 2.39 | 2.32 | 0.83 | 0.98 |
| | Glu | 6.50 | 6.63 | 0.97 | 0.98 |

**B. TR and non-TR patients (excluding those with short follow-ups); metabolite levels are normalized by tCr.**

| Brain region | Metabolite | mean (TR) | mean (non-TR) | p-value | q-value |
|---|---|---|---|---|---|
| ACC | **GSH** | 0.24 | 0.26 | **6.68E-03** | 0.13 |
| | NAA | 1.22 | 1.24 | 0.42 | 0.78 |
| | GABA | 0.29 | 0.29 | 0.74 | 0.82 |
| | Glu | 1.40 | 1.38 | 0.74 | 0.82 |
| CSO | GSH | 0.23 | 0.23 | 0.62 | 0.78 |
| | NAA | 1.42 | 1.39 | 0.28 | 0.78 |
| | GABA | 0.29 | 0.31 | 0.08 | 0.78 |
| | Glu | 1.03 | 1.04 | 0.61 | 0.78 |
| DLPFC | GSH | 0.27 | 0.26 | 0.85 | 0.90 |
| | NAA | 1.46 | 1.46 | 0.96 | 0.96 |
| | GABA | 0.29 | 0.27 | 0.20 | 0.78 |
| | Glu | 1.25 | 1.28 | 0.28 | 0.78 |
| OFR | GSH | 0.26 | 0.27 | 0.38 | 0.78 |
| | NAA | 1.39 | 1.39 | 0.56 | 0.78 |
| | GABA | 0.29 | 0.31 | 0.40 | 0.78 |
| | Glu | 1.15 | 1.20 | 0.14 | 0.78 |
| Thal | GSH | 0.25 | 0.24 | 0.62 | 0.78 |
| | NAA | 1.30 | 1.32 | 0.49 | 0.78 |
| | GABA | 0.43 | 0.41 | 0.52 | 0.78 |
| | Glu | 1.14 | 1.14 | 0.62 | 0.78 |

**C. TR and all non-TR patients (including those with short follow-ups); metabolite levels are normalized by water.**

| Brain region | Metabolite | mean (TR) | mean (non-TR) | p-value | q-value |
|---|---|---|---|---|---|
| ACC | **GSH** | 1.40 | 1.49 | **5.75E-03** | 0.11 |
| | NAA | 7.06 | 7.14 | 0.21 | 0.73 |
| | GABA | 1.71 | 1.68 | 1.00 | 1.00 |
| | Glu | 8.06 | 7.93 | 0.47 | 0.89 |
| CSO | GSH | 1.29 | 1.32 | 0.81 | 0.95 |
| | NAA | 8.32 | 8.21 | 0.31 | 0.78 |
| | GABA | 1.68 | 1.80 | 0.07 | 0.67 |
| | Glu | 5.99 | 6.12 | 0.25 | 0.73 |
| DLPFC | GSH | 1.45 | 1.53 | 0.61 | 0.89 |
| | NAA | 7.98 | 8.32 | 0.14 | 0.73 |
| | GABA | 1.59 | 1.52 | 0.67 | 0.89 |
| | Glu | 6.83 | 7.18 | 0.21 | 0.73 |
| OFR | GSH | 1.48 | 1.51 | 0.98 | 1.00 |
| | NAA | 7.87 | 7.61 | 0.49 | 0.89 |
| | GABA | 1.67 | 1.65 | 0.71 | 0.89 |
| | Glu | 6.53 | 6.58 | 0.68 | 0.89 |
| Thal | GSH | 1.43 | 1.43 | 0.93 | 1.00 |
| | NAA | 7.43 | 7.50 | 0.59 | 0.89 |
| | GABA | 2.39 | 2.25 | 0.26 | 0.73 |
| | Glu | 6.50 | 6.50 | 0.54 | 0.89 |

**D. TR and all non-TR patients (including those with short follow-ups); metabolite levels are normalized by tCr.**

| Brain region | Metabolite | mean (TR) | mean (non-TR) | p-value | q-value |
|---|---|---|---|---|---|
| ACC | **GSH** | 0.24 | 0.26 | **2.32E-03** | **4.64E-02** |
| | NAA | 1.22 | 1.25 | 0.09 | 0.73 |
| | GABA | 0.29 | 0.30 | 0.68 | 0.91 |
| | Glu | 1.40 | 1.38 | 0.92 | 0.92 |
| CSO | GSH | 0.23 | 0.23 | 0.75 | 0.91 |
| | NAA | 1.42 | 1.41 | 0.63 | 0.91 |
| | GABA | 0.29 | 0.31 | 0.11 | 0.73 |
| | Glu | 1.03 | 1.05 | 0.45 | 0.91 |
| DLPFC | GSH | 0.27 | 0.27 | 0.68 | 0.91 |
| | NAA | 1.46 | 1.46 | 0.78 | 0.91 |
| | GABA | 0.29 | 0.26 | 0.18 | 0.73 |
| | Glu | 1.25 | 1.26 | 0.34 | 0.90 |
| OFR | GSH | 0.26 | 0.27 | 0.33 | 0.90 |
| | NAA | 1.39 | 1.37 | 0.57 | 0.91 |
| | GABA | 0.29 | 0.30 | 0.71 | 0.91 |
| | Glu | 1.15 | 1.19 | 0.18 | 0.73 |
| Thal | GSH | 0.25 | 0.25 | 0.61 | 0.91 |
| | NAA | 1.30 | 1.32 | 0.36 | 0.90 |
| | GABA | 0.43 | 0.40 | 0.86 | 0.91 |
| | Glu | 1.14 | 1.13 | 0.84 | 0.91 |

#### TABLE S6. Longitudinal analysis results for brain volume.

Significant results (q-value < 0.05) are in bold with a gray shadow. Abbreviations: APC represents annual percentage change.

| Metabolite | Average APC (%) | Bayes factor | q-value (T test) |
|---|---|---|---|
| right hippocampus | -0.08 | 0.17 | 0.77 |
| left superior frontal gyrus | 0.28 | 0.25 | 0.47 |
| left gyrus rectus | -1.45 | **18.69** | **0.01** |
| left middle occipital gyrus | 1.29 | **7.30** | **0.01** |

**TABLE S7. The performance of classification models**

We compared the performance of five classification models: 1) model of age of onset (denoted as Onset); 2) model of glutathione in anterior cingulate cortex (denoted as GSH); 3) model of right hippocampal volume (denoted as HP); 4) model of left superior frontal gyral volume (denoted as SFG); and 5) model of multimodal markers (denoted as Onset + GSH + HP + SFG). Abbreviations: AUC represents area under the curve; CI, confidence interval; and AIC, Akaike information criterion.

**A. TR and non-TR patients (excluding those with short follow-ups)**

| Predictor | AUC | CI | AIC |
|---|---|---|---|
| Onset | 0.65 | 0.48-0.82 | 53.20 |
| GSH | 0.70 | 0.53-0.87 | 51.17 |
| HP | 0.56 | 0.38-0.74 | 56.36 |
| SFG | 0.58 | 0.40-0.76 | 56.27 |
| Onset + GSH + HP + SFG | 0.84 | 0.71-0.97 | 37.59 |

**B. TR and non-TR patients (including those with short follow-ups)**

| Predictor | AUC | CI | AIC |
|---|---|---|---|
| Onset | 0.64 | 0.49-0.79 | 78.06 |
| GSH | 0.63 | 0.47-0.79 | 79.68 |
| HP | 0.61 | 0.45-0.77 | 79.53 |
| SFG | 0.58 | 0.42-0.74 | 81.35 |
| Onset + GSH + HP + SFG | 0.80 | 0.67-0.93 | 63.53 |

### SUPPLEMENTARY FIGURE LEGENDS

**FIGURE S1. Classification models of treatment resistant (TR) and non-TR patients (including those with short follow-ups).**

We compared the performance of five classification models: 1) model of age of onset (denoted as Onset), blue, AUC=0.64; 2) model of glutathione in anterior cingulate cortex (denoted as GSH), purple line, AUC=0.63; 3) model of right hippocampal volume (denoted as HP), green line, AUC=0.61; 4) model of left superior frontal gyral volume (denoted as SFG), orange line, AUC=0.58; and 5) model of multimodal markers (denoted as Onset + GSH + HP + SFG), red line, AUC=0.80.

**FIGURE S2. Future perspective.**

We have established a multimodal cohort of first episode psychosis patients and healthy controls with a collection of clinical and neuropsychological test scores, molecular measures from different types of biospecimens, and neuroimaging data. Multimodal cohorts like ours could provide a comprehensive understanding of diseases through hypothesis and/or data-driven approaches. We expect that such cohorts will become essential resources for tackling the diagnosis and treatment of brain disorders.

**Figure S1.**

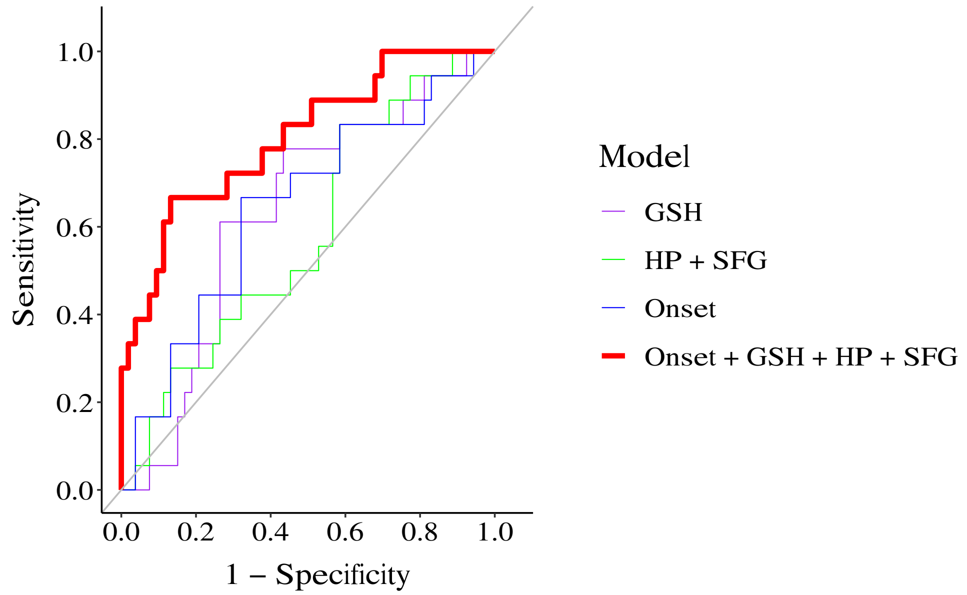
1.0
0.8
0.6
0.4
0.2
0.0
Sensitivity
0.0
0.2
0.4
0.6
0.8
1.0
1 − Specificity
Model
GSH
HP + SFG
Onset
Onset + GSH + HP + SFG

**Figure S2.**

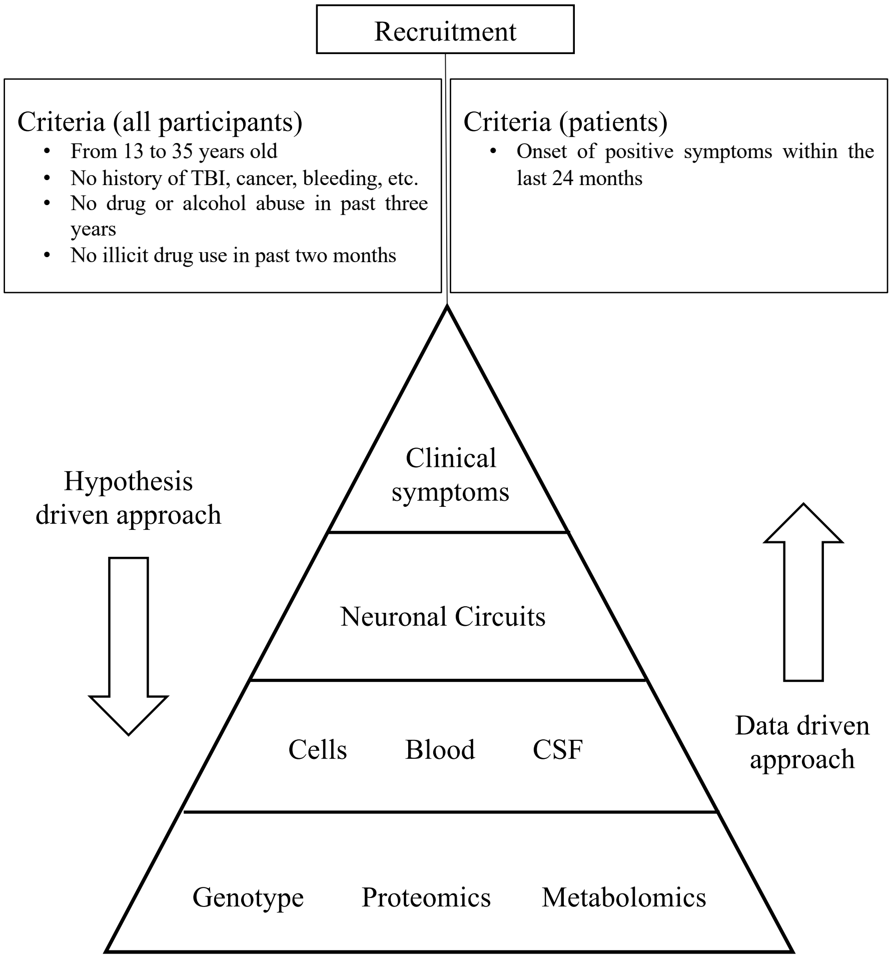
Recruitment
Criteria (all participants)
• From 13 to 35 years old
• No history of TBI, cancer, bleeding, etc.
• No drug or alcohol abuse in past three years
• No illicit drug use in past two months
Criteria (patients)
• Onset of positive symptoms within the last 24 months
Hypothesis driven approach
Clinical symptoms
Neuronal Circuits
Cells
Blood
CSF
Genotype
Proteomics
Metabolomics
Data driven approach
